# Identification of genetic factors controlling phosphorus utilization efficiency in wheat by genome-wide association study with principal component analysis

**DOI:** 10.1101/2020.09.04.283556

**Authors:** Luqman Bin Safdar, Muhammad Jawad Umer, Fakhrah Almas, Siraj Uddin, Qurra-tul-Ain Safdar, Kevin Blighe, Umar Masood Quraishi

**Affiliations:** Department of Plant Sciences, Quaid-i-Azam University, Islamabad 45320, Pakistan; Zhengzhou Fruit Research Institute, Chinese Academy of Agricultural Sciences, Zhengzhou, China; Plant Breeding Institute, Faculty of Agriculture & Environment, University of Sydney, Australia; Government College for Women, 266 RB Khurianwala, Faisalabad 37630, Pakistan; Institute of Ophthalmology, 11-43 Bath Street, University College London, London, UK

**Keywords:** P utilization efficiency, Genome-wide association study, Phosphate rock

## Abstract

Despite the economic importance of P utilization efficiency, information on genetic factors underlying this trait remains elusive. To address that, we performed a genome-wide association study in a spring wheat diversity panel ranging from landraces to elite varieties. We evaluated the phenotype variation for P utilization efficiency in controlled conditions and genotype variation using wheat 90K SNP array. Phenotype variables were transformed into a smaller set of uncorrelated principal components that captured the most important variation data. We identified two significant loci associated with both P utilization efficiency and the 1st principal component on chromosomes 3A and 4A: *qPE1-3A* and *qPE2-4A*. Annotation of genes at these loci revealed 53 wheat genes, among which 6 were identified in significantly enriched pathways. The expression pattern of these 6 genes indicated that *TraesCS4A02G481800*, involved in pyruvate metabolism and TCA cycle, had a significantly higher expression in the P efficient variety under limited P conditions. Further characterization of these loci and candidate genes can help stimulate P utilization efficiency in wheat.

**KEY MESSAGE:** We report two new loci for P utilization efficiency on chromosomes 3A and 4A of wheat. The prioritized candidate genes at these loci can be investigated by molecular biology techniques to improve P efficiency in wheat and grass relatives.

## INTRODUCTION

Phosphorus (P) is an essential component of ATP, phospholipids, and nucleic acids, and, therefore, crucial for the structural integrity and regulation of plant metabolism (Vance 2011). Plants uptake P from soil in the form of phosphate ions through the process of diffusion, and, due to the fact that the P diffusion rate is slow (10^−12^-10^−15^ m^2^ s^−1^), it limits plant P uptake (Schachtman et al. 1998). Thus, there must be readily available P in soil to maintain a reliable amount of uptake during the crop cycle. However, the available data suggests that nearly 5.7 billion hectares of land has insufficient available P for sustainable crop production (Fageria et al. 2017). To ensure proper P supply, excessive fertilizer application is often used in agricultural practices. According to estimates from earlier this decade, an annual increase of 2% P fertilizers is needed to maintain proper crop production (Heffer and Prud’homme 2014). However, excessive P fertilizer application has disadvantages. Most of the P in soil is not available for plant uptake because of a high P fixation rate, and, consequently, it leaches down into the deeper soil layers, ultimately contaminating the environment (Fageria et al. 2017). Add to that the fact that the only natural resource of P is the non-renewable rock phosphate, which is rapidly depleting due to the high consumption rates – its reserves are estimated to be completely drained by the end of this century (Cordell et al. 2009). Researchers have also predicted that the P fertilizer prices (390 USD per metric tonne in 2019 (www.indexmundi.com)) may increase in the coming years and, thus, the majority of farmers worldwide will face greater difficulty in affording P fertilizers (Wissuwa et al. 2015). From these facts, improving P efficiency of crops so that the productivity is not affected by a lower supply becomes an important biological question. Another concern is that P uptake alone cannot improve crop P use efficiency, as higher P uptake itself leads to the environmental concerns of P mining in low input cropping systems, and also economic concerns of higher P fertilizer demand in high input cropping systems.

Thus, there is a need to focus on the improvement of P utilization efficiency, which is a complex trait that complements P uptake and that ultimately leads to better grain yield (Rose and Wissuwa 2012). Previous studies have reported a substantial phenotypic variation of P utilization efficiency in various genotypes of the same species (Ozturk et al. 2005; Yuan et al. 2017), which suggests that identifying the genetic factors underlying this variation can be successfully used to improve crop P utilization efficiency.

To identify the genetic variation causing phenotypic variability, genome-wide association based on linkage disequilibrium in natural populations has been a commonly used approach in recent years (Breseghello and Sorrells 2006; Atwell et al. 2010). GWAS has been successfully used to identify many causal genetic variants for human and plant diseases and important agronomic traits including P efficiency, such as *GmACP1* in soybean (Zhang et al. 2014). This study of Zhang *et al.* followed up GWAS with candidate gene association to reveal that six natural polymorphisms in *GmACP1* controlled 33% phenotypic variability for soybean P efficiency. This suggests that GWAS can be used as an extremely useful tool to identify variants for P utilization efficiency in other important crops as well, such as wheat, rice, and maize.

Wheat is an important staple food crop and feeds nearly one-third of the global population according to FAO statistics; however, the genetic mechanism of P utilization efficiency in wheat remains unclear. Therefore, we designed this study to identify the genetic factors controlling the phenotypic variability of P utilization efficiency in a historical spring wheat panel from Pakistan – wheat is the staple food crop in Pakistan. Although P deficiency at all stages affects plant growth, the most drastic reduction in grain yield due to P deficiency stress occurs in the first 3-6 weeks (Gericke 1924; Gericke 1925). Therefore, we conducted a 21-day experiment in controlled lab conditions using a cigar roll method to check the phenotypic variation of seedling traits, P uptake and P utilization efficiency. For the genomic variation, we used a wheat 90K SNP data published in our earlier study for this diversity panel (Safdar et al. 2020). To correlate the phenotypic and genotypic variation, we followed a method recently proposed by Yano *et al.*, where they performed principal component analysis for the phenotypic data to bring the correlated variable into a smaller number of uncorrelated principal components (PCs), which captured the most important information from the original data (Yano et al. 2019). The seedling variables in our data were also correlated, which is why we chose this approach. In our results, we identified two loci on chromosome 3A and 4A that were significantly associated with PC1, PC2 and P utilization efficiency.

Based on linkage disequilibrium between neighboring SNPs, we extracted 66 high confidence protein coding genes from the wheat reference genome RefSeq v1.0 (Appels et al. 2018), among which 53 could be annotated based on the available information from wheat and relative grass species. The metabolic pathway enrichment analysis of these genes indicated two genes to be involved in amino acid and cellular respiration pathways, in which P is a key component. The expression pattern of these two genes (especially *TraesCS4A02G481800*) were validated via RT-qPCR and showed consistency with the phenotype variation for P utilization efficiency. Future research should study the molecular mechanism of these candidate genes identified in this study, in particular the two genes involved in P related metabolic pathways.

## MATERIALS AND METHODS

### Plant material and experimental conditions

A spring wheat diversity panel of 150 genotypes including landraces, pre-green revolution, post-green revolution, and elite varieties was used; the details of the panel were published in our earlier paper (Safdar et al. 2020). The phenotypic experiment was carried out using a previously published cigar roll method (Zhu et al. 2005). This experiment was planted in 3 replicates and the growth conditions were the same as published earlier (Safdar et al. 2020), except for P concentration. The experiment was divided into two treatments: normal P supply; and, reduced P supply to evaluate the phenotypic variation under P deficiency stress. Modified Hoagland solutions were applied having 100% P amount (31 ppm) and 50% P amount (15.5 ppm) in normal and stress conditions, respectively (Hoagland and Arnon 1950). Plants were kept in a growth chamber with 10 h light and 18-25°C temperature and harvested after 21 days from germination.

### Phenotype evaluation and P estimation

Germination rate, fresh weight, shoot length, leaf area, and relative water content were measured according to Pask et al. (2012). The relative chlorophyll index was measured from the second leaf after 14 and 21 days from germination using a chlorophyll meter (SPAD-502 Konica Minolta sensing Inc.). Root analysis was performed using GiA Roots software using the user-defined instructions (Galkovskyi et al. 2012).

P concentration was estimated with the molybdovanadate method using a spectrophotometer from oven-dried 0.5 g leaf samples digested in concentrated HNO_3_ (Nagornyy 2013). P utilization efficiency was calculated as a ratio between plant fresh weight and P concentration (Sandaña 2016).

Basic statistics, correlation between variables and principle component analysis were performed using XLSTAT software. Statistically significant differences between treatments was estimated using a paired samples t-test with α = 0.05 as a significant threshold.

### Genotyping and GWAS

Genotypic variation was estimated using a wheat 90K SNP array. Population structure (Q) was estimated using STRUCTURE software, whereas a relative kinship matrix (K) and linkage disequilibrium (LD) were estimated using TASSEL software. The information about genotype, Q and LD has been provided in our previous studies (Ain et al. 2015; Safdar et al. 2020). GWAS was performed for P, P utilization efficiency, PC1 and PC2 using the commonly used mixed linear model in TASSEL software (Yu and Buckler 2006). The regression model correlating the genotype-phenotype data included Q and K as covariates to account for background variation during the association tests (Quraishi et al. 2011). To account for false positives expected from type-I error during the multiple testing, Bonferroni correction was applied to consider an association statistically significant. According to this, any SNP-trait association at or below the P-value of 0.000047 (1/nSNP: 1/20,853) was considered significant.

### Gene annotation and pathway enrichment analysis

Based on pairwise LD between neighboring SNPs, physical genomic intervals were assigned to the identified loci for extracting genes. Wheat genes and annotations were extracted from the gff3 files available at the EnsemblPlants database for wheat reference genome RefSeq v1.0 (ftp://ftp.ensemblgenomes.org/pub/plants/release-47/gff3/triticum_aestivum). Due to the fact that information about the wheat genes is quite limited, information of orthologous genes from rice was also extracted using the BioMart tool in the EnsemblPlants database. To identify the role of these genes in biological pathways, KEGG pathway analysis was performed using the KOBAS online tool (Xie et al. 2011). Orthologous genes with high homology from rice were used for enrichment analysis, as the wheat genome is not available in any of the pathway enrichment databases. Genes identified for metabolic pathways were selected for gene expression analysis.

### Gene expression analysis

Primer markers for selected genes were designed with Primer3 plus and are listed in **Supplemental Table 1**. The transcript abundance was analyzed using RT-qPCR in three independent biological replicates of two wheat varieties. The two varieties were selected for comparison based on their P utilization efficiencies: NARC-09 (23.25%) and MH-97 (5.75%). Samples for RNA extraction were collected on day 14 after germination after 6 h light exposure. RNA was isolated with RNA kit (TIANGEN, Beijing, China) following the set protocols. Similarly, cDNA was also synthesized following the instruction of the manufacturer (Takara, Tokyo, Japan). Reaction details were the same as provided previously (Jawad et al. 2020). The wheat tubulin gene was used as an internal check to normalize the RT-qPCR results (Li et al. 2014).

**Table 1.** Descriptive statistics and analysis of variance for seedling traits and phosphorus utilization efficiency.

| Trait<sup>a</sup> | Control |  |  | P Stress |  |  | Control x P Stress<sup>d</sup> | P Stress Impact<sup>e</sup> |
| --- | --- | --- | --- | --- | --- | --- | --- | --- |
|  | Mean ± SE | VC<sup>b</sup> | SK (P)<sup>c</sup> | Mean ± SE | VC<sup>b</sup> | SK (P)<sup>c</sup> |  |  |
| GR (%) | 85.26 ± 1.29 | 0.15 | -0.67 | 84.57 ± 0.88 | 0.10 | -0.43 | - | -0.81 % |
| Chl<sub>14DAG</sub> | 23.84 ± 0.29 | 0.12 | -0.26 | 24.67 ± 0.26 | 0.11 | -0.55 | * | +3.48 % |
| Chl<sub>21DAG</sub> | 23.56 ± 0.33 | 0.14 | -0.26 | 24.22 ± 0.25 | 0.11 | -0.26 | * | +2.80 % |
| FW (mg) | 114.38 ± 2.25 | 0.20 | 0.38 | 91.52 ± 2.03 | 0.22 | 0.84 | *** | -19.99 % |
| SL (cm) | 17.69 ± 0.33 | 0.19 | 0.63 | 18.03 ± 0.28 | 0.16 | 1.29 | * | +1.92 % |
| LA (cm<sup>2</sup>) | 4.33 ± 0.08 | 0.20 | 1.04 | 3.68 ± 0.07 | 0.20 | 0.73 | *** | -15.01 % |
| RWC | 76.34 ± 0.92 | 0.12 | -0.82 | 74.26 ± 0.62 | 0.08 | -0.36 | * | -2.72 % |
| RL (cm/cm<sup>3</sup>) | 0.03 ± 0.00 | 0.38 | 0.00 | 0.05 ± 0.00 | 0.32 | 0.06 | *** | +66.67 % |
| RW (cm) | 5.16 ± 0.11 | 0.22 | 0.48 | 5.24 ± 0.08 | 0.15 | -0.45 | * | +1.55 % |
| NW:ND | 0.29 ± 0.01 | 0.60 | 1.66 | 0.38 ± 0.01 | 0.43 | 1.60 | *** | +31.03 % |
| P (mg/kg) | 11.14 ± 0.44 | 0.39 | 0.59 | 9.13 ± 0.28 | 0.31 | 0.79 | *** | -18.04 % |
| PUtE (%) | 11.98 ± 0.56 | 0.47 | 1.58 | 10.90 ± 0.41 | 0.38 | 1.73 | ** | -9.02 % |

<sup>a</sup>Germination Rate (GR), Chlorophyll Content 14th Day of Germination (Chl<sub>14DAG</sub>), Chlorophyll Content 21st Day of Germination (Chl<sub>21DAG</sub>), Fresh Weight (FW), Shoot Length (SL), Leaf Area (LA), Relative Water Content (RWC), Primary Root Length (RL), Root Width (RW): this represents the root diameter, computed from the root images for all pixels of the medial axis of root system, Network Width to Depth Ratio (NW:ND): network width is estimated from the images as all pixels from left to right and network depth as all pixels from top to bottom, Phosphorus Concentration in Leaf (P), Phosphorus Utilization Efficiency (PUtE).
<sup>b</sup> Coefficient of variation.
<sup>c</sup>Skewness based on Pearson's principle. Traits that showed skewness were square root transformed to normalize the data before genetic association analysis.
<sup>d</sup> \*, \*\* and \*\*\* represent significant differences at $P \leq 0.05$ , $P \leq 0.01$ and $P \leq 0.001$ , computed using a paired sample t-test.
<sup>e</sup> Impact of P deficiency stress in terms of percent increase or decrease.

## RESULTS

### Phenotypic evaluation and principal component analysis

P deficiency stress severely impacted the normal plant growth and reduced the plant fresh weight by 20%, leaf area by 15%, network area by 13.4%, P concentration by 18% and P utilization efficiency by 9%. Root length increased significantly under stress (+66%) and the chlorophyll content also showed a marginal increase both after 14 and 21 days from germination (**Table 1**). Trait correlations were all statistically significantly positive, except for root length with P concentration in control treatment (*r*^2^ = −0.23) and P stress treatment (*r*^2^ = −0.31), and between P concentration and P utilization efficiency in control treatment (*r*^2^ = −0.77) and P stress treatment (*r*^2^ = −0.64). These correlations were understandable because, when the P supply is not enough, roots tend to elongate in search of P (Gahoonia and Nielsen 2003); also, when a plant uptakes excess P, it cannot be utilized sufficiently by the plant Detailed correlation results, box plots, and QQ scatter plots are presented in **Figure 1**.

**Figure 1.**
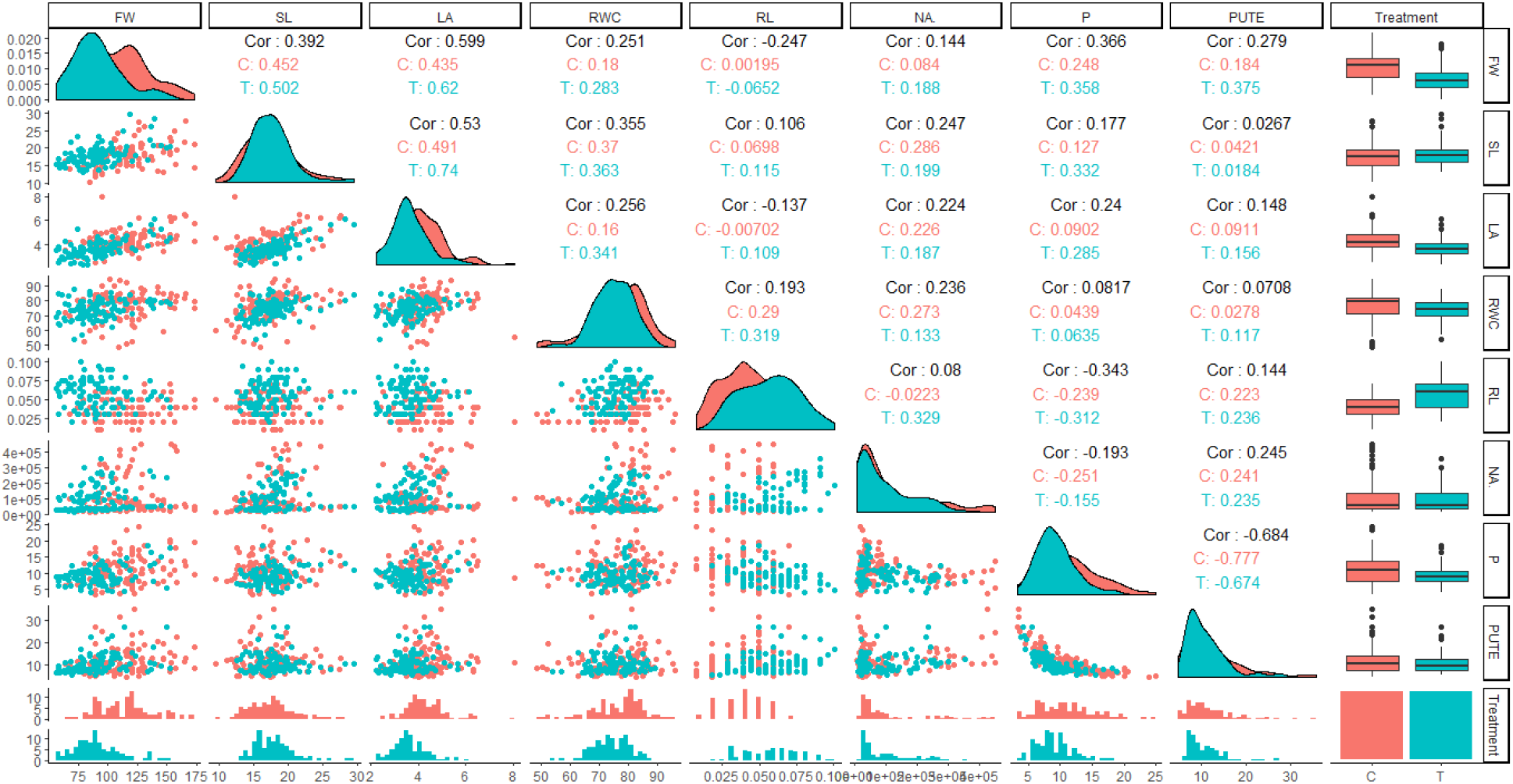
Phenotype distribution of seedling traits and coefficient of correlation. The bottom panel and the center line cutting the figure into two triangles represent histograms for each trait. The right panel represent box and whiskers for trait distribution. The lower triangle represents QQ scatter plots for trait distribution. The upper triangle represents coefficient of correlation (*r*^2^ values) between variables. Axes of histograms and scatter plots represent numerical values for phenotype. **\*cf.** Trait nomenclature is presented in **Table 1** legends.

To identify the factors underlying phenotypic variation, we performed principal component analysis for all traits. For normal P treatment, all traits showed positive loadings on PC1, however, P concentration showed a much higher loading (0.9) on PC2. PC1 and PC2 explained more than 50% of the total phenotypic variation (**Figure 2a-c**). For P stress conditions, P utilization efficiency (0.85) and root length (0.54) showed highly positive loadings on PC2, while morphological traits again showed highly positive loadings on PC1. P concentration also showed positive loading (0.54) on PC1. Together, PC1 and PC2 explained more than 60% phenotypic variation from P deficiency stress treatment (**Figure 2d-f**). These results suggest that PC1 can be used as a quantitative index for P concentration and morphological traits, and PC2 for P utilization efficiency under stress conditions. Therefore, we used PC1 and PC2 transformed from stress variables for GWAS analysis. We also included P concentration and P utilization efficiency in GWAS analysis to ensure that any useful information is not missed by PCs.

**Figure 2.**
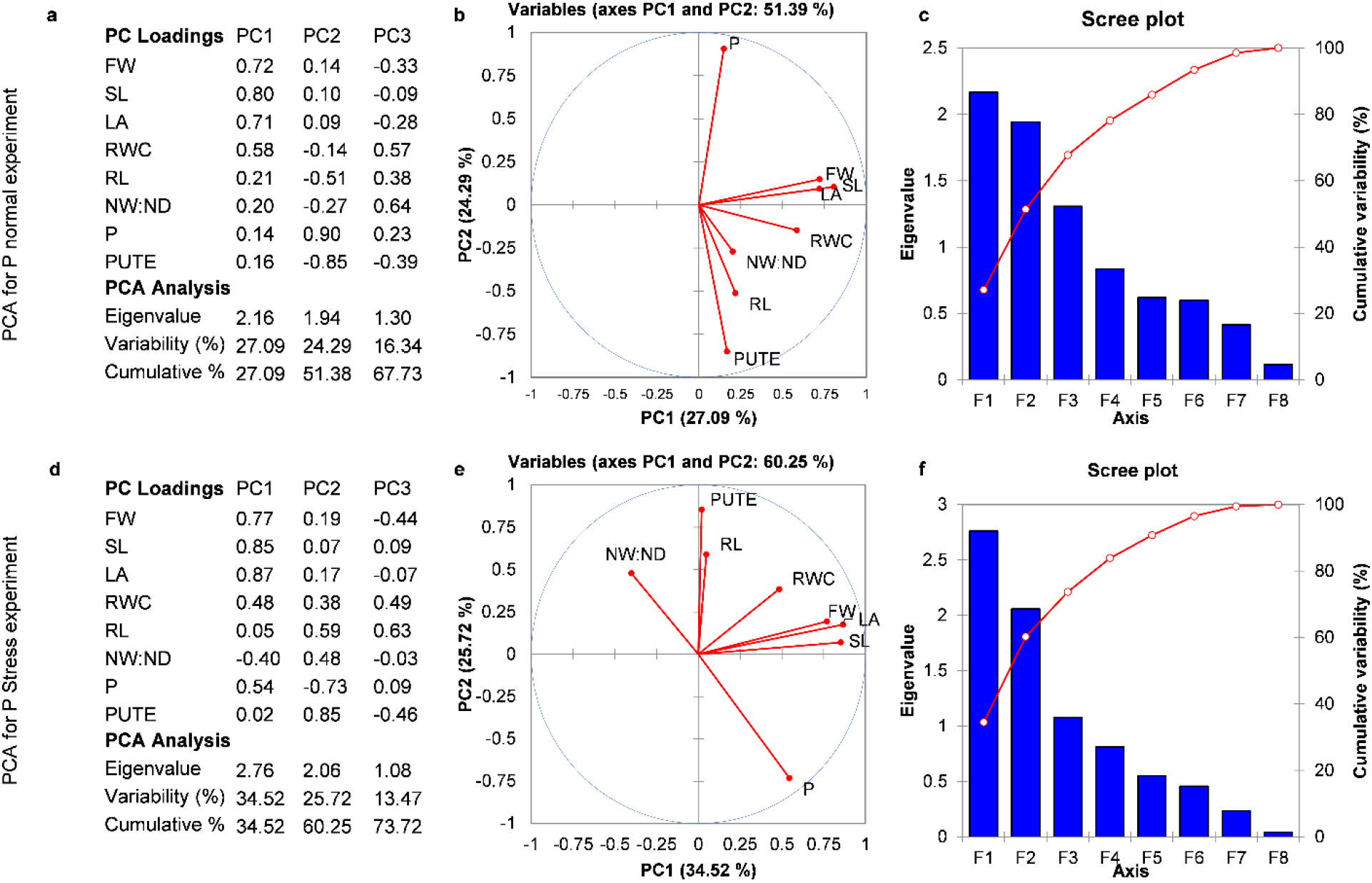
PCA for trait variables: **a-c)** represent PCA for normal P conditions data; **d-f)** represent PCA for limited P conditions data. **a)** PC loadings for trait variables in first three PCs and variability % explained by them under normal P. All traits show positive loadings on PC1 but the loadings for root and P related traits are comparatively much lower. **b)** A 2D PCA plot showing the distribution of various traits on PC1 and PC2. Together, first two PCs explain more than 50% phenotype variation from the data under normal growth conditions. **c)** Scree plot showing eigenvalues for all the PCs and the variability explained by individual PC. **d)** PC loadings for trait variables in first three PCs and variability % explained by them under P stress. P utilization efficiency and root length show highly positive loadings on PC2. **b)** A 2D PCA plot showing the distribution of various traits on PC1 and PC2. Together, first two PCs explain more than 60% phenotype variation from the data under P stress. **c)** Scree plot showing eigenvalues for all the PCs and the variability explained by individual PC. The eigenvalues for first two PCs and their difference from the rest are higher for P stress data compared to the normal treatment data. **\*cf.** Trait nomenclature is presented in **Table 1** legends.

### Genotype dataset and population structure

There were originally 81,587 markers in the 90K Infinium array, which were filtered to remove the monomorphic markers, those with sequencing errors and with a minor allele frequency less than 5%. Finally, 20,853 high quality polygenic SNP markers were kept for the GWAS analysis. Marker density was highest in the B sub-genome, followed by A and D. Population structure, which divided the panel into seven sub-groups, was used to account for the background variation. Population linkage disequilibrium decay was 800 kb in sub-genome B, 500 kb in sub-genome D, and 300 kb in sub-genome A. Details about each dataset, population structure, and linkage disequilibrium are reported in our earlier studies (Ain et al. 2015; Safdar et al. 2020) - readers are encouraged to see those papers for further details.

### SNP-trait associations from GWAS analysis

From the genotype-phenotype association tests for PC1, PC2, P concentration and P utilization efficiency from P stress treatment data, five statistically significant SNP-trait associations were identified in total (**Table 2**). Two consistent association peaks appeared for P utilization efficiency on chromosomes 3A and 4A, with one SNP at each peak reaching up to the statistical significance (**Figure 3a**). Three SNPs significantly associated with P concentration on chromosomes 1A, 3D, and 7A (**Figure 3b**). The same peak and SNP that associated with P utilization efficiency at chromosome 4A showed a significant association with PC1 (**Figure 3c**). For PC2, both regions that associated with P utilization efficiency showed upward peaks but no SNP could pass statistical significance (**Figure 3d**). These results, collectively, suggested that the two loci identified at chromosomes 3A and 4A were important for P utilization efficiency, with 4A locus more likely being the causal loci as it associated with both P utilization efficiency and PC1. These two loci were named as *qPE1-3A* and *qPE2-4A* and further investigated for the identification of candidate genes for P utilization efficiency. The SNP at *qPE1-3A* was IWB38606, with a physical position of 3A:12980739 on the wheat reference genome, while the SNP at *qPE2-4A* was IWB59368, with a physical position of 4A:737340127.

**Table 2.** Five significant loci identified by genome-wide association study.

| Loci<sup>a</sup> | SNP ID<sup>b</sup> | Chr | Position<sup>c</sup> | P-value<sup>d</sup> | Phenotype | Treatment |
| --- | --- | --- | --- | --- | --- | --- |
| *qPU1-1A* | IWA1376 | 1A | 1177154 | 4.50E-05 | P uptake | P stress |
| *qPE1-3A* | IWB38606 | 3A | 12980739 | 2.98E-05 | PUtE | P stress |
| *qPU2-3D* | IWB13582 | 3D | 604304010 | 3.62E-05 | P uptake | P stress |
| *qPE2-4A* | IWB59368 | 4A | 737340127 | 4.39E-05 | PC1 | P stress |
| *qPE2-4A* | IWB59368 | 4A | 737340127 | 4.39E-05 | PUtE | P stress |
| *qPU3-7A* | IWB72514 | 7A | 1739310 | 3.88E-05 | P uptake | P stress |

<sup>a</sup> Loci are named based on their association with phenotype in this study.
<sup>b</sup> Universal SNP IDs used for wheat 90K Infinium SNP genotype array.
<sup>c</sup> Physical position of SNP on wheat reference genome IWGSC RefSeq v1.0.
<sup>d</sup> Significant p-values calculated by Bonferroni multiple testing correction.

**Figure 3.**
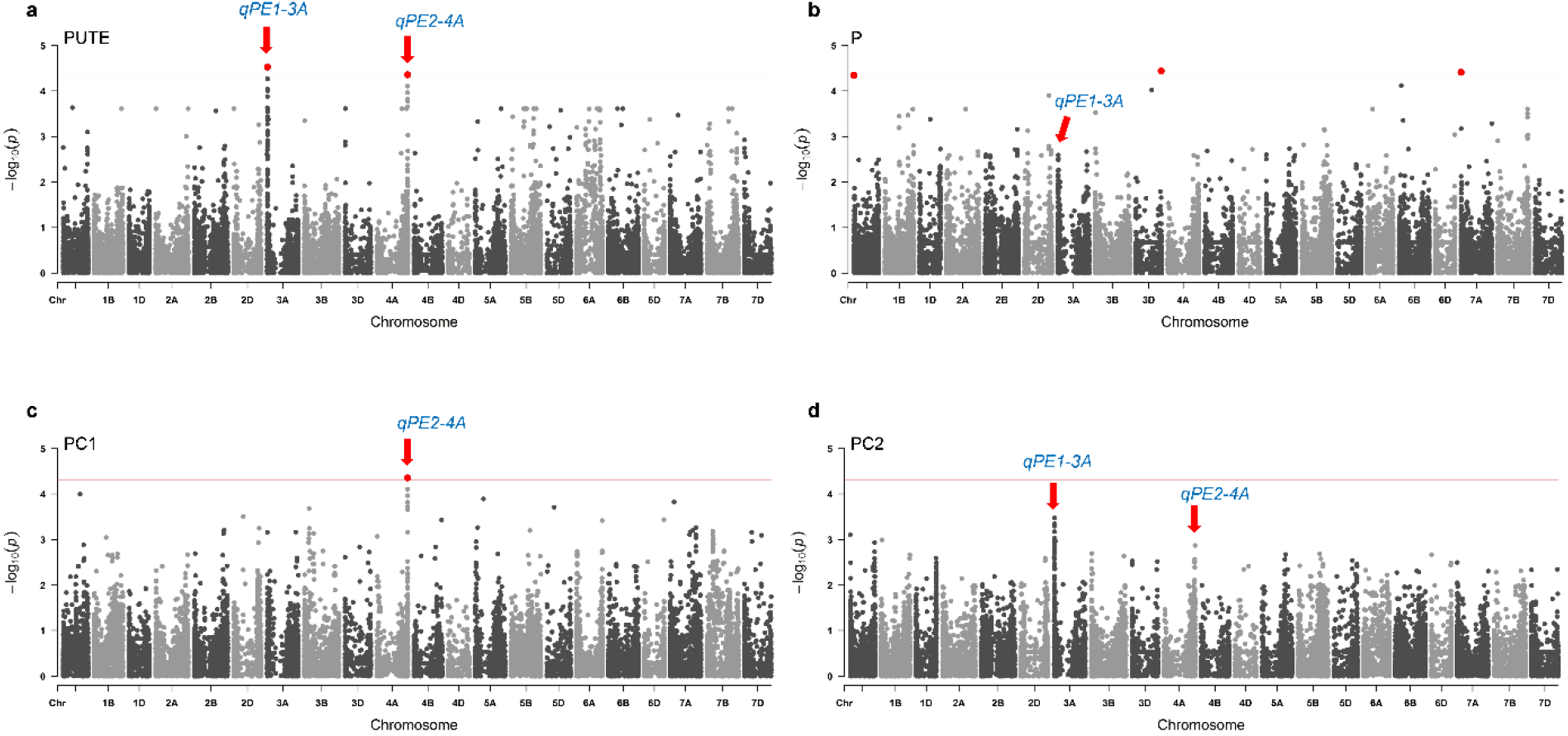
Association tests for P utilization efficiency, P uptake, PC1 and PC2. The red line in the plots shows the statistical significance after applying correction for multiple comparisons. **a)** P utilization efficiency significantly associates at two position: chromosome 3A and 4A. Both peaks are consistent and one SNP at each locus is statistically significant. Significant SNPs are colored red. **b)** Three SNPs are identified statistically significant for P uptake, but no consistent peaks are observed for these SNPs. **c)** Test result of PC1 shows the similar significant association observed for P utilization efficiency at chromosome 4A. **d)** No statistically significant association is observed for PC2, however, the same loci at 3A and 4A show somewhat weak correlation.

### Genes identified at two P utilization efficiency loci

For the identification of candidate genes, we estimated the pairwise LD between the neighboring SNPs at the two significant loci (*qPE1-3A* and *qPE2-4A*). Based on pairwise LD, the two regions were allocated 260 kb (*qPE1-3A*) and 120 kb (*qPE2-4A*) intervals (**Figure 4**). The number of genes extracted from the wheat reference genome were 75 at 3A locus and 58 at 4A locus (**Supplemental Table 2**). Among these were 66 high confidence protein coding genes: 40 at *qPE1-3A* and 26 at *qPE2-4A* (**Supplemental Table 3**). As the information about wheat gene annotation is quite limited, we also extracted gene annotation for orthologous *Oryza sativa* Japonica rice genes since both crops belong to the same family. Together, these provided the annotation information of 53 genes (**Table 3**); thus, the other 13 genes for which no annotation was available were excluded from further analysis.

**Table 3.** Wheat genes present at two PUtE associated loci (*qPE1-3A* and *qPE2-4A*) and their rice homologs with available descriptors from wheat and rice databases from EnsemblPlant

| Wheat Gene ID | Rice Gene ID | Gene description | %ID |
| --- | --- | --- | --- |
| *TraesCS4A02G481800* | *Os06g0105400* | Acetyltransferase component of pyruvate dehydrogenase complex | 75.64 |
| *TraesCS3A02G025200* | *Os01g0121900* | E3 ubiquitin-protein ligase | 39.53 |
| *TraesCS3A02G025500* | *Os01g0120800* | Eukaryotic translation initiation factor 3 subunit A | 78.80 |
| *TraesCS3A02G022600* | *Os08g0465800* | Glutamate decarboxylase | 78.87 |
| *TraesCS3A02G023600* | *Os06g0190800* | Glutamate receptor | 63.66 |
| *TraesCS3A02G023900* | *Os05g0102400* | Protein NEN4 | 60.62 |
| *TraesCS4A02G481400* | *Os06g0105600* | Protein transport protein sec16 | 67.27 |
| *TraesCS3A02G025700* | *Os01g0121100* | Protein-lysine N-methyltransferase | 80.41 |
| *TraesCS3A02G024400* | *Os04g0112300* | tRNA (adenine(58)-N(1))-methyltransferase non-catalytic subunit TRM6 | 74.95 |
| *TraesCS3A02G022500* | *Os02g0496500* | HhH-GPD domain containing protein | 47.39 |
| *TraesCS3A02G022700* | *Os11g0621300* | Protein of unknown function DUF1399 family | 62.58 |
| *TraesCS3A02G022800* | *Os03g0235100* | Similar to Pg4 | 83.74 |
| *TraesCS3A02G022900* | *Os01g0119000* | Glycosyltransferase AER61 | 59.13 |
| *TraesCS3A02G023000* | *Os01g0119000* | Glycosyltransferase AER61 | 61.00 |
| *TraesCS3A02G023100* | *Os01g0119100* | Similar to Glycosyltransferase | 65.58 |
| *TraesCS3A02G023200* | *Os12g0641100* | Similar to Na<sup>+</sup>/H<sup>+</sup> antiporter | 79.79 |
| *TraesCS3A02G023400* | *Os01g0152800* | Protein kinase, core domain containing protein | 54.90 |
| *TraesCS3A02G023800* | *Os01g0120300* | Similar to EMB2369 (EMBRYO DEFECTIVE 2369); ATP binding / aminoacyl-tRNA ligase/ leucine-tRNA ligase/ nucleotide binding | 88.67 |
| *TraesCS3A02G024100* | *Os01g0120400* | Alanyl-tRNA synthetase, class IIc family protein | 72.73 |
| *TraesCS3A02G024200* | *Os01g0120500* | Similar to PAP fibrillin domain containing protein | 72.05 |
| *TraesCS3A02G024300* | *Os01g0120500* | Similar to PAP fibrillin domain containing protein | 74.02 |
| *TraesCS3A02G024500* | *Os05g0542900* | Pectin lyase fold domain containing protein | 61.35 |
| *TraesCS3A02G024600* | *Os01g0120600* | Similar to Oxidoreductase NAD-binding domain containing protein | 69.15 |
| *TraesCS3A02G024700* | *Os07g0574600* | Hypothetical conserved gene | 22.06 |
| *TraesCS3A02G024900* | *Os01g0120700* | Similar to sarcoplasmic reticulum histidine-rich calcium-binding protein | 64.21 |
| *TraesCS3A02G025100* | *Os03g0176100* | Similar to predicted protein | 49.65 |
| *TraesCS3A02G025300* | *Os07g0586100* | Similar to Esterase precursor (EC 3.1.1.-) (Early nodule-specific protein homolog) | 70.37 |
| *TraesCS3A02G025400* | *Os07g0586100* | Similar to Esterase precursor (EC 3.1.1.-) (Early nodule-specific protein homolog) | 72.84 |
| *TraesCS3A02G025600* | *Os01g0121000* | Conserved hypothetical protein | 57.71 |
| *TraesCS3A02G025800* | *Os01g0121200* | Zinc finger, RING/FYVE/PHD-type domain | 78.07 |
| *TraesCS3A02G025900* | *Os05g0211100* | Similar to Cytochrome P450-like protein | 67.60 |
| *TraesCS3A02G026000* | *Os01g0121300* | Conserved hypothetical protein | 74.81 |
| *TraesCS3A02G026200* | *Os01g0121600* | Conserved hypothetical protein | 61.90 |
| *TraesCS3A02G026400* | *Os01g0121600* | Conserved hypothetical protein | 59.18 |
| *TraesCS3A02G026500* | *Os01g0121700* | ABC transporter-like domain containing protein | 89.37 |
| *TraesCS4A02G481700* | *Os06g0105500* | Protein arginine N-methyltransferase domain | 78.19 |
| *TraesCS4A02G482000* | *Os06g0105350* | GRAS (GAI-RGA-SCR) plant-specific transcription factor, Maintenance of shoot apical meristem indeterminacy | 70.00 |
| *TraesCS4A02G482100* | *Os05g0528900* | Ribosomal protein L9 family protein | 78.64 |
| *TraesCS4A02G482300* | *Os11g0689300* | Similar to Calmodulin-binding protein | 74.46 |
| *TraesCS4A02G482500* | *Os02g0227500* | Protein kinase, catalytic domain containing protein | 34.24 |
| *TraesCS4A02G482600* | *Os02g0227500* | Protein kinase, catalytic domain containing protein | 33.42 |
| *TraesCS4A02G482700* | *Os02g0227500* | Protein kinase, catalytic domain containing protein | 36.68 |
| *TraesCS4A02G482800* | *Os11g0567800* | Leucine-rich repeat-containing N-terminal | 36.49 |
| *TraesCS4A02G482900* | *Os02g0227500* | Protein kinase, catalytic domain containing protein | 35.05 |
| *TraesCS4A02G483000* | *Os02g0227500* | Protein kinase, catalytic domain containing protein | 33.15 |
| *TraesCS4A02G483200* | *Os11g0567800* | Leucine-rich repeat-containing N-terminal | 41.89 |
| *TraesCS4A02G483300* | *Os02g0227500* | Protein kinase, catalytic domain containing protein | 36.68 |
| *TraesCS4A02G483400* | *Os10g0375400* | Leucine-rich repeat domain containing protein | 51.63 |
| *TraesCS4A02G483500* | *Os11g0567800* | Leucine-rich repeat-containing N-terminal | 14.86 |
| *TraesCS4A02G483700* | *Os11g0558400* | Leucine-rich repeat, N-terminal domain | 38.99 |
| *TraesCS4A02G483900* | *Os02g0227500* | Protein kinase, catalytic domain containing protein | 35.60 |
| *TraesCS4A02G484000* | *Os11g0524601* | Similar to NB-ARC domain containing protein | 17.33 |

%ID. *Triticum aestivum* gene identical to target *Oryza sativa* Japonica Group gene

**Figure 4.**
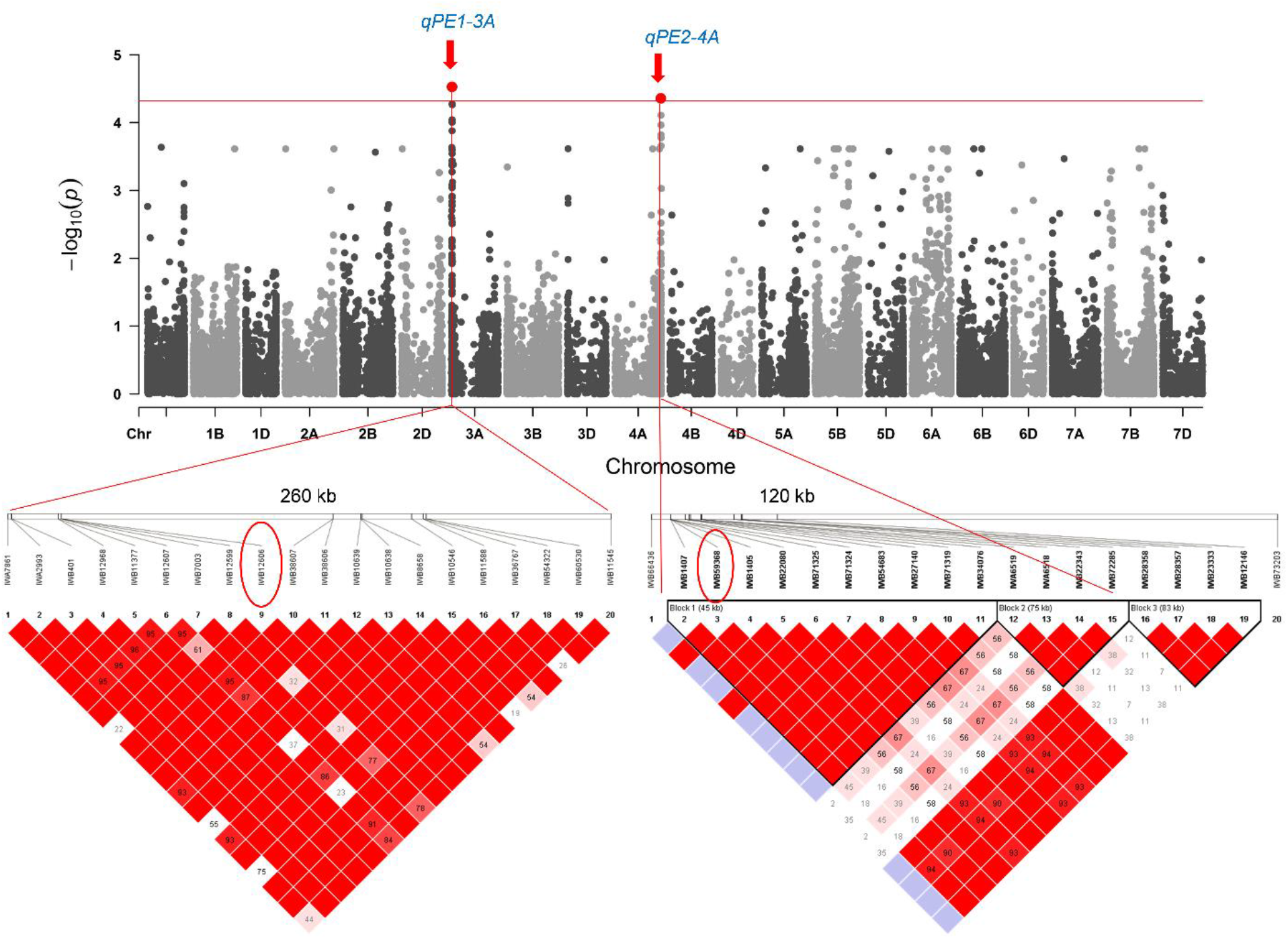
Genomic length of two significant SNP loci based on pairwise LD between neighboring SNPs. The first locus at 3A covers 260 kb distance, whereas the second locus at 4A covers 120 kb distance. The LD decay distance for sub-genome A in this mapping population is 300 kb.

### Genes identified in the metabolic pathways

The 53 genes identified at *qPE1-3A* and *qPE2-4A* were analyzed for pathway enrichment using the KEGG database. Six out of these 53 genes were involved in the top 10 identified pathways and among them two genes were identified in more than one pathway (**Table 4**). One of these two genes was present at 3A (*TraesCS3A02G022600*) involved in taurine and hypotaurine metabolism, butanoate metabolism, and steroids biosynthesis. The second gene was identified at 4A (*TraesCS4A02G481800*) and is involved in TCA cycle and pyruvate metabolism, both of which are interconnected with P utilization efficiency. From these results, *TraesCS4A02G481800* looked like a strong candidate for P utilization efficiency as the locus *qPE2-4A* was also identified as the more likely causal locus (significantly associated with both P utilization efficiency and PC1). However, we performed reverse transcription quantitative PCR for all these six genes to see their expression pattern before suggesting any of these genes as potential candidates.

**Table 4.** Candidate genes from significant loci involved in top ten KEGG pathways.

| Wheat Gene<sup>a</sup> | Rice Homolog | KEGG Pathway<sup>b</sup> | BN<sup>c</sup> |
| --- | --- | --- | --- |
| *TraesCS3A02G022600* | *Os08g0465800* | Taurine and hypotaurine | 21 |
| *TraesCS3A02G022600* | *Os08g0465800* | Butanoate metabolism | 27 |
| *TraesCS3A02G025900* | *Os05g0211100* | Steroid biosynthesis | 32 |
| *TraesCS3A02G022600* | *Os08g0465800* | beta-Alanine metabolism | 46 |
| *TraesCS3A02G025100* | *Os03g0176100* | Basal transcription factors | 47 |
| *TraesCS3A02G022600* | *Os08g0465800* | Aspartate and glutamate metabolism | 52 |
| *TraesCS4A02G481800* | *Os06g0105400* | Citrate cycle (TCA cycle) | 58 |
| *TraesCS4A02G481800* | *Os06g0105400* | Pyruvate metabolism | 86 |
| *TraesCS3A02G023800* | *Os01g0120300* | Aminoacyl-tRNA biosynthesis | 115 |
| *TraesCS3A02G025200* | *Os01g0121900* | Ubiquitin mediated proteolysis | 136 |

<sup>a</sup> Genes from *qPE1-3A* and *qPE2-4A* in top metabolic pathways.
<sup>b</sup> Metabolic pathway from KEGG database of homologous rice genes.
<sup>c</sup> Background number of identified pathways.

### Expression profiles of the six genes identified in metabolic pathways

The expression pattern of the six genes was evaluated in two varieties: MH-97 with low P utilization efficiency (5.75%); and NARC-09 with high P utilization efficiency (23.25%). RNA extracted after 14 days from germination was used for gene expression analysis since the early stage P deficiency is known to cause severe damage to plant growth even when the supply becomes reliable at the later stages (Gericke 1924; Gericke 1925). All six genes showed a significantly higher relative expression in both varieties under normal P conditions. Only two genes (*TraesCS3A02G022600*, *TraesCS4A02G481800*) showed a difference in expression between the two varieties. *TraesCS4A02G481800* showed a much prominent and statistically significant difference in expression and up-regulation in P efficient variety NARC-09 under stress conditions (**Figure 5**). These results again indicate that these two genes were involved in the P utilization efficiency of plants, and, between these two, *TraesCS4A02G481800* (present at *qPE2-4A*) had a considerably higher contribution. This further suggests that the 4A locus controlled the phenotypic variation for P utilization efficiency. This locus can be thoroughly investigated for P utilization efficiency in future studies, although the candidate genes identified here also appear to be potentially causal genes.

**Figure 5.**
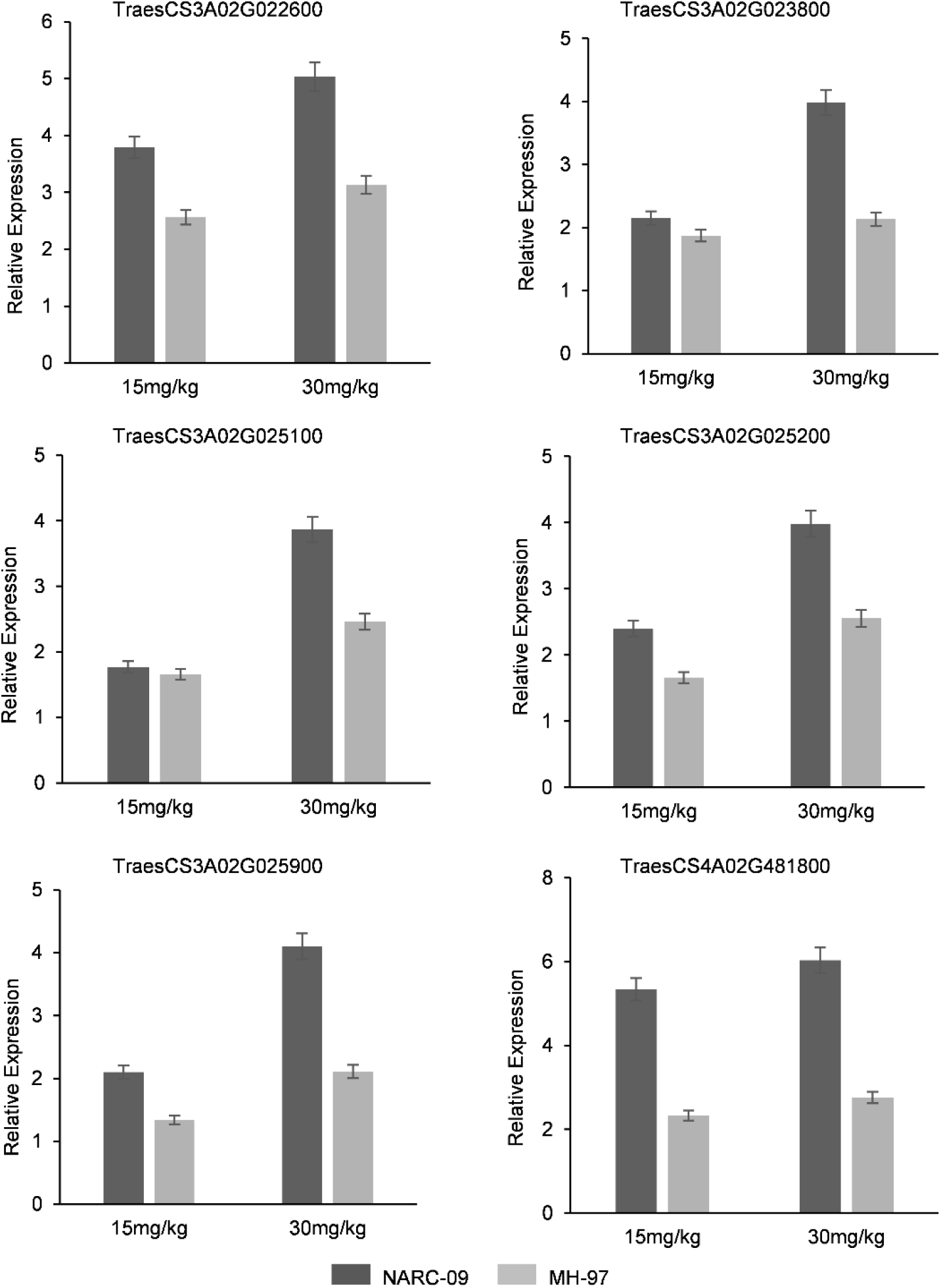
The expression pattern of six candidate genes that are identified in the top ten KEGG pathways based on information for homologous genes in rice. The only 4A locus gene among these six shows more than two-folds difference and higher expression in P efficient variety. MH-97 (P deficient) and NARC-09 (P efficient) were selected based on P utilization efficiency data.

## DISCUSSION

In recent years, a large number of studies have established that P deficiency in agricultural soils and the continuously depleting natural P reserves will be a grave threat to the optimal yield of cereal crops in future. To improve the overall P efficiency of crops, it is essential to ensure that plants maximize their use of the P that they uptake from soil. However, despite the significance of this trait, no information is available about the genetic factors underlying P utilization efficiency in wheat. In the present study, we applied a genome-wide association model to identify variants for P utilization efficiency.

The germplasm we used in this study consisted of spring wheat varieties from Pakistan with enormous genetic diversity. It included landraces, varieties released before the green revolution, varieties released after the green revolution and elite varieties released in the 21st century. Such diversity in genetic material is key to explore the factors underlying complex traits like P efficiency. Nearly a decade ago, phosphorus starvation tolerance gene (*PSTOL1*) was characterized in rice by Gamuyao *et al*. When they overexpressed this gene in P inefficient rice varieties, the grain yield and biomass increased under P limited growth conditions. Their study showed that *PSTOL1* acted as an enhancer of early root growth resulting in a higher P uptake in plants. An interesting observation of that study was that *PSTOL1* gene was not present in the rice reference genome, which indicated the significance and need of exploring traditional germplasm resources to dissect complex traits (Gamuyao et al. 2012). Apart from being historically diverse, the germplasm from Pakistan also confers value because most of the agricultural soils are calcareous in nature, where P uptake is often limited due to a high soil P fixation rate (Friesen et al. 1997), and as a result, there is a substantial phenotypic variation for P uptake and utilization efficiency within the panel. Therefore, the variants identified from this germplasm could be of great significance to identify causal genes for P utilization efficiency in wheat.

Evolutionarily, rice and wheat are close relatives, both belonging to grass family *Gramineae*, and many orthologous genes in one species are often found to perform the identical functions in the other species. Similarly, for P efficiency, efforts were made to improve P use efficiency in wheat based on the information from rice. Milner *et al.* recently cloned *TaPSTOL* gene (an orthologous gene of rice *PSTOL1*) and investigated its effects on wheat under various P supply conditions (Milner et al. 2018). They observed that the expression of *TaPSTOL* was higher in root tips and hairs under limited P supply and the overexpression or silencing significantly impacted agronomic traits. However, this impact was not correlated with P supply conditions, which made it difficult to conclude whether or not this gene contributed to P efficiency in wheat. Another fact is that wheat is a hexaploid crop and the evolution of polyploid crop species has been a rather intriguing series of events: gene retention/loss during their evolution has been asymmetrical, resulting in the events of duplication, sub-functionalization, and neofunctionalization (Liu et al. 2014). Therefore, it is not necessarily true that homologous genes should always perform similar functions in related species. Hence, the information about the genetic control of P utilization in wheat remains largely unknown and the results of the present study may substantiate the genomic selection for yield maximization in P stress conditions.

In this study, we used principal components from phenotype data to identify genetic variants due to the fact that this approach had recently identified a causal gene for plant architecture in rice (Yano et al. 2019). The two significant loci that we identified for P utilization efficiency under stress were also explained by the first two principal components, especially *qPE2-4A*. Pathway enrichment analysis of the genes in these two loci (*qPE1-3A* and *qPE2-4A*) narrowed our focus to six genes, which were identified in top ten metabolic pathways (**Table 4**). We further observed that *TraesCS4A02G481800* was involved in pyruvate metabolism and TCA cycle. These two are interconnected reactions and central to cellular respiration, without which cells cannot produce energy necessary for proper functioning, ultimately leading to death. The role of inorganic P is crucial in this reaction because a phosphate group replaces coenzyme A from succinyl, forming a high-energy bond. When succinyl converts into succinate, this energy is utilized in substrate-level phosphorylation to form guanine triphosphate (GTP), which can be converted into ATP (Martínez-Reyes and Chandel 2020). Together, all these facts indicate that *TraesCS4A02G481800* is involved in P efficiency and could be considered as a potential candidate. The transcript abundance analysis further indicated that this gene had a significantly higher expression in P efficient wheat variety under 50% P supply conditions. However, conclusive inference should not be drawn before validating the phenotype by transgene studies and understanding the complete characteristic mechanism of this gene alone and in relation to the gene-network.

The loci identified in our study did not colocalize with the previously reported loci for P use efficiency or the loci identified for P concentration in this study. The understandable reason behind this dysconnectivity is that previous studies have mainly focused on P uptake efficiency of crops. However, physiological studies have reported that P uptake alone cannot enhance the overall P efficiency of the plant (Rose and Wissuwa 2012) due to the fact that the utilization of the absorbed P also varies greatly across individual genotypes. Therefore, P utilization efficiency is critical in terms of ensuring sustainable production in the future. Our results further support this phenomenon, as the identified SNPs for P concentration and P utilization efficiency were present at separate loci and linkage groups on the genome. These results, however, signify the objective of studying the genetic basis of P utilization efficiency.

One limitation of our study should also be noted: although the natural population selected for this study was genetically diverse and incorporated a potentially wide range of alleles from landrace to elite cultivars, the individual sample sizes within the 7 subpopulations were relatively smaller. A larger diversity panel including samples from geographically and historically diverse populations can present interesting information on rare alleles underlying P utilization efficiency that may have been selected against during the course of domestication and breeding improvement, as this trait has not been targeted in breeding programs.

Future molecular biology studies should investigate the effects of these candidate genes on the agronomic traits of wheat in P stress conditions. The two newly identified potential loci can also be explored further using comparative genomics approaches. In conclusion, these results add significantly to our existing knowledge of P utilization efficiency in bread wheat.

## Acknowledgements

The authors thank Dr. Muhammad Rizwan Alam (Dept. of Biochemistry, Quaid-i-Azam University) for helping in P estimation.

## Funding statement

This study was supported by the funding from Higher Education Commision, Pakistan, Grant/Award: HEC NRPU-3825 to Umar Masood Quraishi.

## Author contributions

UMQ designed the study and provided the financial and experimental resources. LBS, MJU and FA performed the phenotyping and experimental work. LBS performed the statistical/bioinformatics analysis and wrote the manuscript. SU and QS helped in the data analysis. KB and UMQ edited and revised the manuscript.

## Compliance to ethical standards

All research was conducted following the research safety standards of Pakistan Agriculture Research Council and Quaid-i-Azam University.

## Conflict of interest

The authors declare that the research was conducted in the absence of any commercial or financial relationships that could be construed as a potential conflict of interest.

## Supplementary information

**Supplementary Table 1.** Marker sequences used for gene amplification in RT-qPCR.

**Supplementary Table 2.** A complete list genes present at two significant GWAS loci.

**Supplementary Table 3.** High confidence protein coding genes at two significant GWAS loci.

## Notes

### Competing Interest Statement

The authors have declared no competing interest.

